# Subjective cognitive decline predicts lower cingulo-opercular network functional connectivity in individuals with lower neurite density in the forceps minor

**DOI:** 10.1101/2022.03.25.485749

**Authors:** Adriana L. Ruiz-Rizzo, Raymond P. Viviano, Ana M. Daugherty, Kathrin Finke, Hermann J. Müller, Jessica S. Damoiseaux

**Affiliations:** Department of Psychology, General and Experimental Psychology Unit, LMU Munich, Munich, 80802, Germany; Department of Psychology and Institute of Gerontology, Wayne State University, Detroit, MI, 48202, U.S.A.; Department of Neurology, Jena University Hospital, Jena, 07747, Germany

**Keywords:** functional connectivity, latent growth curve modeling, neurite density, salience network, subjective cognitive decline

## Abstract

Cognitive complaints of attention/concentration problems are highly frequent in older adults with subjective cognitive decline (SCD). Functional connectivity in the cingulo-opercular network (CON-FC) supports cognitive control, tonic alertness, and visual processing speed. Thus, those complaints in SCD may reflect a decrease in CON-FC. Frontal white-matter tracts such as the forceps minor exhibit age- and SCD-related alterations and, therefore, might influence the CON-FC decrease in SCD. Here, we aimed to determine whether SCD predicts an impairment in CON-FC and whether neurite density in the forceps minor modulates that effect. To do so, we integrated cross-sectional and longitudinal analyses of multimodal data in a latent growth curve modeling approach. Sixty-nine healthy older adults (13 males; 68.33 ± 7.95 years old) underwent resting-state functional and diffusion-weighted magnetic resonance imaging, and the degree of SCD was assessed at baseline with the memory functioning questionnaire (greater score indicating more SCD). Forty-nine of the participants were further enrolled in two follow-ups, each about 18 months apart. Baseline SCD did not predict CON-FC after three years or its rate of change (*p*-values > 0.092). Notably, however, the forceps minor neurite density did modulate the relation between SCD and CON-FC (intercept; *b* = 0.21, 95% confidence interval, CI, [0.03, 0.39], *p* = 0.021), so that SCD predicted a greater CON-FC decrease in older adults with relatively lower neurite density in the forceps minor. The neurite density of the forceps minor, in turn, negatively correlated with age. These results suggest that CON-FC alterations in SCD are dependent upon the forceps minor neurite density. Accordingly, these results imply modifiable age-related factors that could help delay or mitigate both age and SCD-related effects on brain connectivity.

## Introduction

Subjective cognitive decline (SCD) is the perceived worsening of cognitive abilities, unrelated to an acute event and characterized by normal performance in standard neuropsychological tests (Molinuevo et al., 2017). SCD is associated with changes in the functional connectivity (FC – or the observed correlations between distant regions) (Friston, 1994) within and between the default mode network and medial temporal lobe regions (Viviano & Damoiseaux, 2020). Less is known about SCD-related differences in other potentially relevant networks such as the ‘cingulo-opercular’ network (CON), in which FC decreases in aging and mild cognitive impairment (He et al., 2014). CON-FC supports cognitive control, tonic alertness, and visual processing speed (Dosenbach et al., 2007; Ruiz-Rizzo et al., 2018; Sadaghiani & Kleinschmidt, 2016) that are all vulnerable to decline in aging (e.g., McAvinue et al.,2012; Ruiz-Rizzo et al., 2019). As cognitive complaints related to attention/concentration problems are highly frequent in older adults with SCD (La Joie et al., 2016; Valech et al., 2018), CON-FC might also be associated with SCD. Furthermore, the pattern of observed differences in FC has been proposed to vary according to the time since SCD onset (e.g., increases at the start of SCD and decreases if approaching a mild cognitive impairment stage; Viviano & Damoiseaux,2020). Accordingly, only with a longitudinal approach can we set an individual’s putative SCD onset and baseline FC. Then, FC changes can more clearly be determined. In the present study, we evaluated SCD-related differences in CON-FC integrating both cross-sectional and longitudinal analyses in a latent growth curve modeling approach.

Although strong FC can exist between brain regions with no direct white matter connection (i.e., there is no exact, one-to-one correspondence between FC and white-matter pathways), structural connections do constrain and provide a global ‘physical substrate’ to FC (Honey et al., 2009; Raichle, 2015; Z. Wang et al., 2015). For example, deep neural networks trained with structural connectivity data can accurately predict FC both at both the group and individual levels, indicative of structure-function coupling in the human connectome (Sarwar et al., 2021). Thus, the white matter in tracts connecting some of the CON regions (i.e., structural connectivity, SC – or the modeled diffusion direction of water molecules; Assaf et al., 2019) might modulate the magnitude of SCD-related differences in CON-FC. Particularly for CON, FC-guided tractography has shown that its SC mainly connects the anterior cingulate cortex with other CON regions (e.g., middle frontal gyrus, anterior insula; Figley et al.,2015). Both reduced CON-SC (i.e., lower fractional anisotropy; Figley et al., 2015) and CON-FC (e.g., Onoda et al., 2012; Ruiz-Rizzo et al., 2019) are observed with more advanced age. The forceps minor, a major commissural tract formed from the projections from the genu of the corpus callosum (Wakana et al., 2004), connects the medial (i.e., anterior cingulate cortex) and lateral surfaces of the frontal lobes and can be identified in humans in vivo with diffusion-weighted imaging (Abe et al., 2004). Thus, we studied the forceps minor as a candidate *proxy* of CON-SC for three main reasons. First, as a *major* white-matter tract, it can be more easily identified in different samples (as compared to smaller, exploratory, sample-specific white-matter tracts), and no assumption about a unitary function-to-structure regional correspondence is required. Second, it is a major frontal structural pathway that has been shown to connect core regions in the CON. And third, its diffusion properties can be measured in vivo.

The particular relevance of the forceps minor white matter in SCD is illustrated by a recent study which reported that fractional anisotropy and mean diffusivity of the forceps minor exhibit alterations in SCD and mild cognitive impairment (e.g., Luo et al., 2020). In addition, lower fractional anisotropy and higher mean diffusivity of commissural tracts, including the forceps minor, have been observed with increasing age (de Groot et al., 2015). The degree of fractional anisotropy in the forceps minor relates to performance in executive function (Ohlhauser et al., 2019) and visual attention (Tu et al., 2018) tasks in SCD and/or mild cognitive impairment – i.e., cognitive functions in which complaints are frequent in SCD (Valech et al., 2018). Lastly, in carriers of autosomal Alzheimer’s disease mutations, increased mean diffusivity in the forceps minor appears to precede symptom onset (Araque Caballero et al., 2018). Accordingly, we hypothesized that individual differences in diffusion metrics of the forceps minor modify the relationship between SCD and CON-FC. We focused on the neurite density of the forceps minor, in particular, because neurite density has been proposed to be a more sensitive and specific marker of pathology than other white-matter diffusion metrics to which it contributes (e.g., fractional anisotropy) – as different combinations of neurite density and orientation dispersion can yield a particular value of fractional anisotropy (Zhang et al., 2012).

In the present study, we aimed to evaluate SCD-related differences in CON-FC and whether CON-SC modulates that effect, by using longitudinal, multimodal data. More specifically, we analyzed CON-FC and CON-SC across three time points, ~ 18 months apart, in a sample of healthy older adults in which the degree of SCD was quantified at baseline. We used latent growth curve modeling to provide a flexible estimation of variability *between* individuals in the patterns of change *within* individuals (Curran et al., 2010), which is particularly relevant when not all individuals follow the same trajectory or exhibit the same rate of change, as is the case in aging. Notably, growth modeling is flexible regarding accommodating complex patterns of missing data or compound-shaped trajectories (Curran et al., 2010), which is common in longitudinal data as well as in aging-related processes. Moreover, in the framework of structural equation modeling, latent factors that represent *unobserved* growth trajectories can also be incorporated in the growth model (Curran et al., 2010). As, in this framework, variables can be simultaneously studied as ‘independent’ and ‘dependent’, questions related to antecedents and consequences can be adequately addressed (Duncan & Duncan, 2009). For these reasons we used latent growth curve modeling to test the following two hypotheses: (i) baseline SCD predicts a decrease in CON-FC over three years, and (ii) the decrease in CON-FC is more pronounced in older adults with relatively lower CON-SC as obtained from the neurite density in the forceps minor.

## Materials and Method

### Participants

Sixty-nine healthy older participants (13 males; mean age: 68.33 ± 7.95 years; age range: 50 - 85 years; 78% self-identified as Black) underwent neuropsychological testing and resting-state functional and diffusion-weighted magnetic resonance imaging (MRI) at baseline. Participants were volunteers recruited from the community of Metro Detroit (MI, USA). Forty-nine of those 69 participants were enrolled in a longitudinal study, which consisted of two follow-ups, each approximately 18 months apart (mean: 18.5 ± 1.39 months). Thirty-four (69%) of the 49 participants returned for the first follow-up and 29 (55%) for the second follow-up. CON-SC data were missing for 8 participants at baseline and 1 participant at the last follow-up, and MFQ was missing for 1 participant at baseline. Little’s MCAR test was not significant (χ^2^ (108, *N* = 69) = 100, *p* = 0.690, missing patterns = 9), supporting the conclusion that data were missing at random. The attrition of CON-FC or CON-SC longitudinal data was not associated with age at baseline, sex, racial ethnicity, SCD status or complaints, education, global cognition, depressive symptoms, or IQ (*F*(9, 54) = 1.15, *p* = 0.343). All participants were right-handed and fulfilled the selection criteria, including no recent or past history of psychiatric disorders, neurological disease, or head trauma, no current use of psychoactive medication, and no contraindications for MRI. The Institutional Review Board of Wayne State University approved all the procedures used in this study, which followed the ethical principles of the World Medical Association Declaration of Helsinki. All participants gave written informed consent before their participation in the study. Demographic data for each measurement occasion are listed in Table 1. The longitudinal functional MRI and the baseline diffusion-weighted MRI data used in the current study have been reported previously (Viviano et al., 2019; Viviano & Damoiseaux, 2021).

**Table 1.**
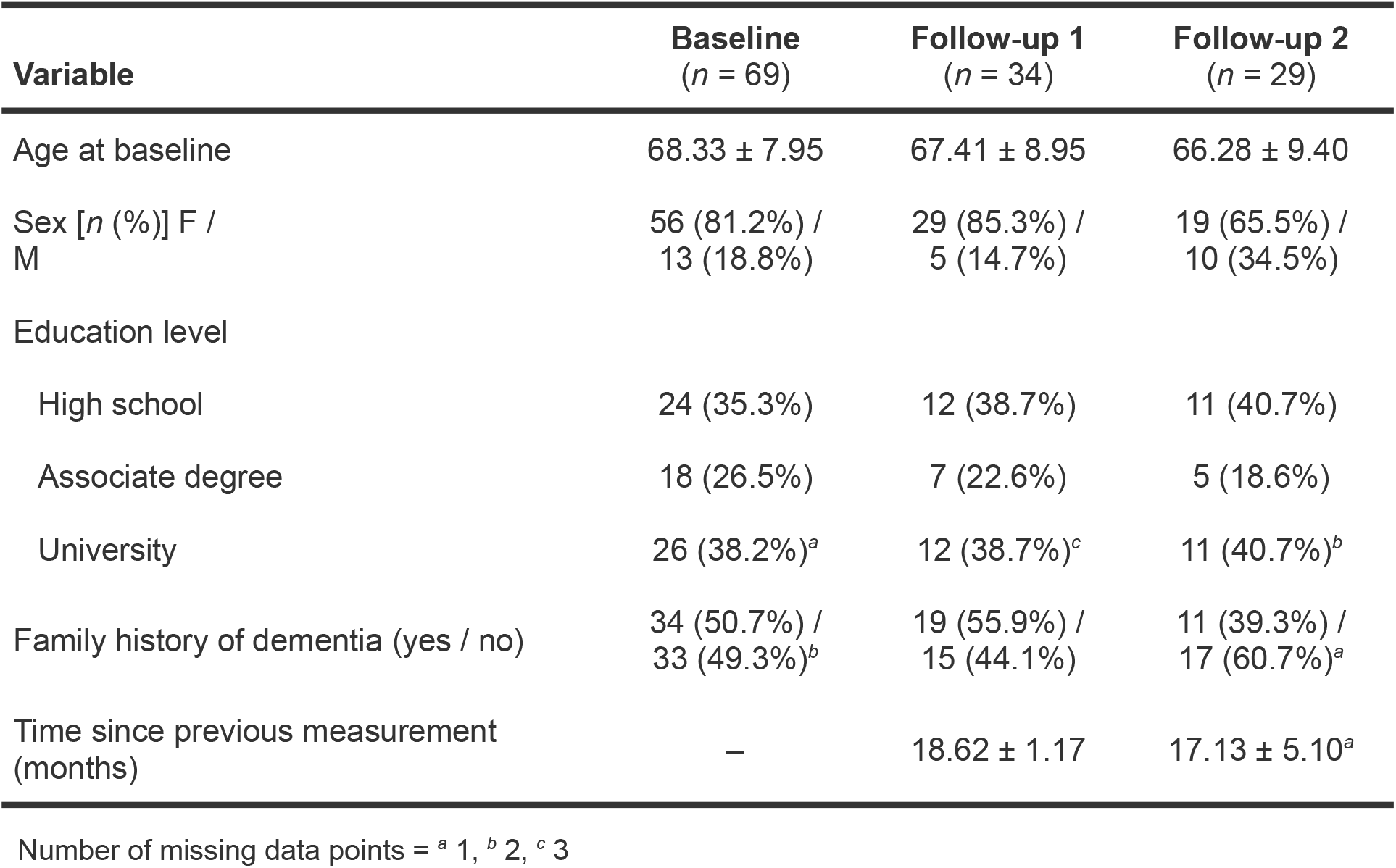
Demographic variables across the three time points.

### SCD, behavioral, and neuropsychological measures

SCD was quantified with the General Frequency of Forgetting factor of the Memory Functioning Questionnaire (MFQ; Gilewski et al., 1990). These scores were inverted^1^ to obtain a more intuitive interpretation of them, with higher scores indicating a greater degree of SCD. Depressive symptoms were measured with the Geriatric Depression Scale (Yesavage et al., 1982), and the Big Five Inventory (John et al., 1991) was used to measure personality traits of neuroticism and conscientiousness. Global cognition was assessed through the Mini-Mental State Examination (MMSE; Folstein et al.,1975) and the Wechsler Abbreviated Scale of Intelligence II (Wechsler, 2011). To assess attention, we used the Trail Making Test (TMT; Reitan & Wolfson, 1986), part A, and the digit symbol-coding subtest of the Wechsler Adult Intelligence Scale III (Wechsler, 1997). The Rey Auditory Verbal Learning Task (Rey, 1958) and the adult battery of the Wechsler Memory Scale-IV (Wechsler, 2009) were used to assess memory. Executive function measures were obtained through the TMT, part B, and the Stroop test (Stroop, 1935). Finally, a measure of language was obtained through a semantic fluency task (animals and occupations). Average group performance in all neuropsychological measures is presented in Table 2, separately for each measurement occasion. To evaluate measurement invariance in neuropsychological assessments across study occasions and SCD, linear mixed-effects models were fitted to the data using the *Ime4* package in R (v. 4.1.0; R Core Team, 2021; RStudio Team, 2021; https://www.R-project.org/). Model parameters were assessed by 95% confidence intervals (CI), which if not overlapping zero support the interpretation of the effect size. Pearson correlations and analysis of variance were used to assess the associations with CON-SC and, respectively, (continuous and categorical) demographic variables.

**Table 2.**
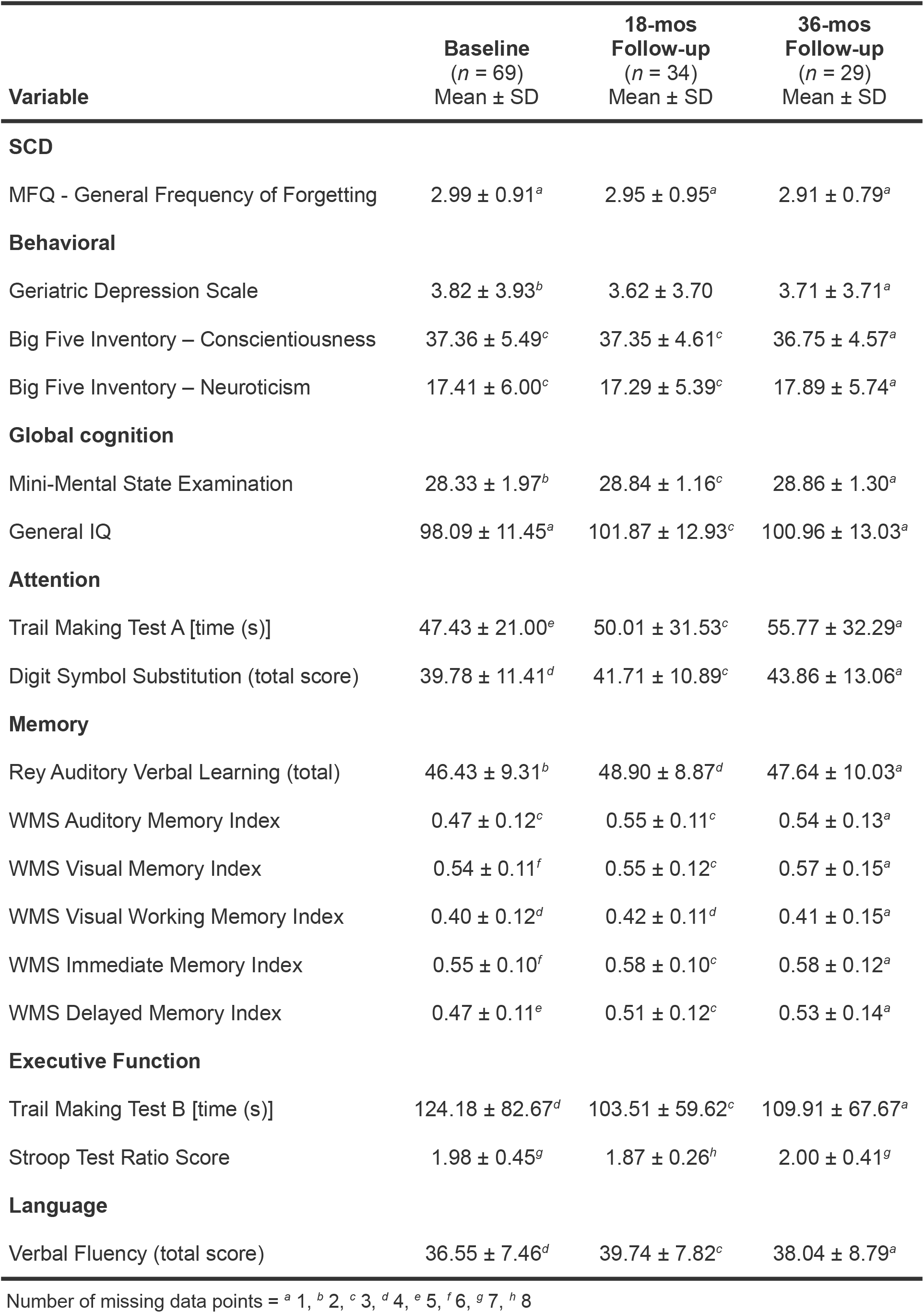
SCD, behavioral, and neuropsychological measures across the three time points

### MRI data acquisition

Structural MRI data were acquired on a Siemens Magnetom Verio 3-Tesla scanner (Siemens Healthcare, Erlangen, Germany) located at the Wayne State University MR Research Facility (Detroit MI, United States), using a 32-channel Head Matrix coil. A high-resolution anatomical image was obtained through a T1-weighted 3D magnetization-prepared rapid gradient-echo (MP-RAGE) sequence, with the following parameters: 176 slices parallel to the bicommissural line, slice thickness = 1.3 mm, repetition time (TR) = 1680 ms, echo time (TE) = 3.51 ms, inversion time = 900 ms, flip angle = 9°, bandwidth = 180 Hz/pixel, GRAPPA acceleration factor = 2, field of view (FOV) = 256 mm, matrix size = 384 × 384, and voxel size = 0.7 × 0.7 × 1.3 mm.

Diffusion-weighted MRI data were acquired through a multiband (acceleration factor = 3) echo-planar imaging (EPI) sequence, with the following parameters: 84 axial slices, slice thickness = 2 mm, TR = 3500 ms, TE = 87 ms, flip angle = 90°, refocus flip angle = 160°, bandwidth = 1724 Hz/pixel, GRAPPA acceleration factor = 2, FOV = 200 mm, matrix size = 100 × 100, voxel size = 2.00 mm isotropic, phase encoding = anterior to posterior, diffusion directions = 96 (6 without diffusion weighting, b = 0), b-values = 1000 (30 directions) & 1800 (60 directions) s/mm^2^, total time of acquisition (TA) = 6 min 1 s.

Resting-state functional MRI data were acquired through a high-resolution multiband (acceleration factor = 3) T2*-weighted EPI sequence, with the following parameters: 75 slices, slice thickness = 2 mm, TR = 2000 ms, TE = 30 ms, flip angle = 73°, bandwidth = 1698 Hz/pixel, GRAPPA acceleration factor = 2, FOV = 256 mm, matrix size = 128 × 128, voxel size = 2 mm isotropic, phase encoding = anterior to posterior, 220 volumes, TA = 7 min 36 s. Participants were asked to remain with their eyes closed during the resting-state functional MRI procedure.

A field map for susceptibility-derived distortion correction was acquired through two multiband (acceleration factor = 3) spin-echo echo-planar images of opposing phase encoding directions (anterior to posterior and posterior to anterior), with the following parameters: 75 slices, slice thickness = 2 mm, TR = 2412 ms, TE = 51 ms, flip angle = 90°, refocus flip angle = 180°, bandwidth = 1698 Hz/pixel, GRAPPA acceleration factor = 2, FOV = 256 mm, matrix size = 128 × 128, and voxel size = 2 mm isotropic.

### Diffusion-weighted MRI data preprocessing

Prior to preprocessing, the high-resolution, T1-weighted images were anatomically segmented with Freesurfer (v.6.0; Fischl, 2012) (http://surfer.nmr.mgh.harvard.edu/). Preprocessing of diffusion-weighted MRI data was done using the TRActs Constrained by UnderLying Anatomy (TRACULA; https://surfer.nmr.mgh.harvard.edu/fswiki/Tracula) tool (Yendiki et al., 2011) in Freesurfer. Preprocessing included eddy-current distortion correction based on FMRIB Software Library (FSL) tools (v.5.0.8; Jenkinson et al., 2012) (https://fsl.fmrib.ox.ac.uk) and head motion correction by registering the diffusion-weighted to the b = 0 images. Next, the b = 0 image was registered to the high-resolution, T1-weighted image by an affine registration method and, subsequently, to the Montreal Neurological Institute (MNI) standard space template. Finally, the anatomical priors for white-matter tracts were computed by incorporating prior knowledge of the pathways from a set of training subjects (Yendiki et al., 2011). All ensuing analyses of these data were conducted in participants’ native (i.e., diffusion-weighted) space.

### Resting-state fMRI data preprocessing

Resting-state fMRI data were preprocessed as described in Viviano & Damoiseaux (2021). Briefly, preprocessing was conducted using FSL FEAT (Woolrich et al., 2001) (https://fsl.fmrib.ox.ac.uk/fsl/fslwiki/FEAT), and included removal of the first five volumes, head motion correction (Jenkinson et al., 2002), removal of non-brain structures (Smith, 2002), susceptibility-derived distortion correction (Smith et al.,2004), co-registration to the anatomical image and, subsequently, to the MNI 2-mm standard space, spatial smoothing (4-mm full width at half maximum), and 4D grand-mean scaling. An independent component analysis (ICA)-based strategy for automatic removal of motion artifacts (ICA-AROMA; Pruim et al., 2015)was used to detect and regress motion artifacts. In short, ICA-AROMA classifies independent components as ‘motion’ based on high-frequency content, correlation with realignment parameters, edge fraction, and the cerebrospinal fluid fraction (Pruim et al., 2015). Finally, the global signal was regressed out from the resting-state functional MRI data^2^.

### Connectivity analysis

#### Structural connectivity

Major white-matter tracts were automatically reconstructed and labeled in participants’ native space using TRACULA, an automated Bayesian global tractography algorithm (Yendiki et al., 2011). TRACULA uses the “ball-and-stick” model of diffusion (Behrens et al., 2007) and anatomical priors for tract reconstruction, both of which yield a posterior probability for each tract given the diffusion-weighted MRI data (Yendiki et al., 2011). More specifically, FSL’s bedpostX (Bayesian Estimation of Diffusion Parameters Obtained using Sampling Techniques) was used to fit the ball-and-stick model to the diffusion-weighted MRI data at every voxel. Next, the probability distributions were generated for each participant and each tract by fitting the shape of each pathway to the results of the ball-and-stick model of diffusion and to the prior knowledge of the pathway anatomy obtained from the manually annotated set of training subjects in the TRACULA atlas (Yendiki et al., 2011).

The posterior probability distribution map for the forceps minor (thresholded to 0.20 of the maximum probability value – default settings) was used as a region of interest (ROI) to obtain its average neurite density. To do so, we used the Neurite Orientation Dispersion and Density Imaging (NODDI) method (Zhang et al., 2012), as implemented in the AMICO (Accelerated Microstructure Imaging via Convex Optimization) python module (Daducci et al., 2015). NODDI follows a tissue model of three microstructural compartments that affect water diffusion differently: intracellular (space bounded by neurite membranes), extracellular (space around the neurites), and CSF (space occupied by cerebrospinal fluid) (Zhang et al., 2012). Neurite density refers to the volume fraction of the intracellular compartment and might be a sensitive marker of pathology (Zhang et al., 2012). After obtaining the voxelwise neurite density map for each participant, the mean neurite density was computed for the forceps minor. The mean neurite density was weighted by each voxel’s posterior probability of tract membership (as in Viviano et al., 2019): voxels toward the center had greater posterior probabilities and voxels farther from the center had lower posterior probabilities. Thus, voxels closer to the boundary of white and gray matter contributed less to the weighted mean. Such an approach reduces the possible contribution of partial volume effects to the results. We used the mean neurite density of the forceps minor as a measure of CON-SC in the present study. A separate mean neurite density was obtained for each time point. These values were then averaged to form a single measure of CON-SC for the latent growth curve modeling analysis to reduce model complexity and the number of parameters to be estimated given our sample size.

#### Functional connectivity

We selected five ROIs of the CON, namely, the anterior cingulate cortex (ACC), left and right anterior insula, and left and right middle frontal gyrus, and obtained the average time course of each ROI using FSL’s *fslmeants* command (also see *Supplementary Material).* These ROIs were taken from a parcellation of the human cerebral cortex that combined information from sharp changes in architecture, function, connectivity, and topography (Glasser et al., 2016). More specifically, each of these ROIs was composed of several multimodal areas (Table 3). These areas were selected based on the descriptions presented in the supplementary neuroanatomical results of Glasser et al. (2016). To create the ROIs for the present study, the selected multimodal areas in MNI space were first extracted from the original map of 360 areas [https://github.com/brainspaces/glasser360/blob/master/glasser360MNI.nii.gz] and binarized (as all voxels had as intensity value the area number). Next, the selected areas were added together, resampled from 1-mm to 2-mm-isotropic voxel size (i.e., the functional images’ voxel size), thresholded (cutoff = 0.3), and binarized. Finally, we extracted the time courses of each ROI by first reslicing each ROI to each participant’s functional space, which gave each voxel of the ROI a different value depending on the warping. Then, a weighted mean time course was computed across all voxels of the ROI, using the ROI voxels’ values (range: 0 – 1) as weights. ROIs’ time courses were obtained for each participant, separately per time point. Pearson’s correlation was first computed between the five ROI’s time series (i.e., 10 pairs), and the *r*-values were then Fisher’s *r*-to-*z* transformed. Finally, the mean CON-FC was calculated from the *z*-values across the 10 ROI pairs.

**Table 3.**
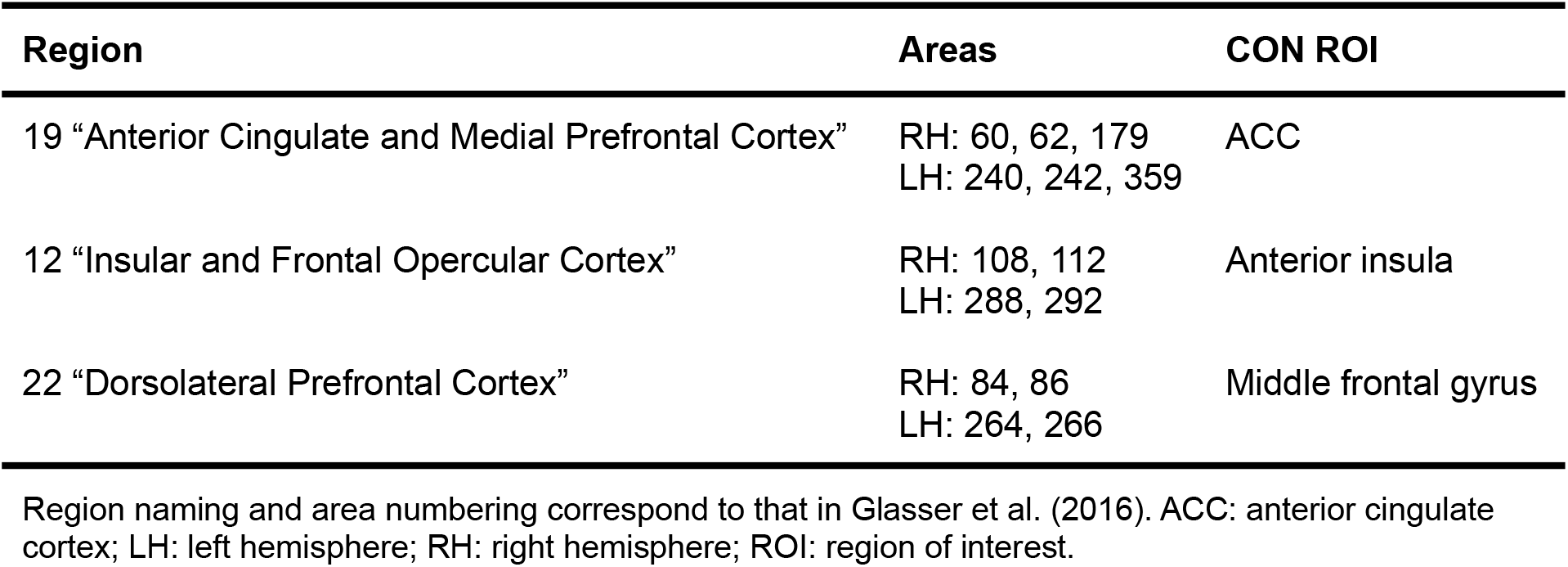
Multimodal areas comprising the CON-FC ROIs

### Latent growth curve modeling

Hypotheses were tested using a latent growth curve model (LGCM; McArdle, 2009) in a longitudinal structural equation model framework. LGCMs allow estimating between-person differences in within-person patterns of change over time (Curran et al., 2010). Specifically, for each individual, the mean and variance of two (unobserved) factors can be obtained, namely, the latent intercept and the latent slope, respectively. The growth model contains both fixed (i.e., mean of all individual trajectories) and random (i.e., variance around the group means) effects for each of these factors (Curran et al., 2010). Accordingly, relevant variables (i.e., those that might be associated with the random components of growth) can also be included in the model with the aim of predicting random variability in the intercept or slope (Curran et al., 2010). One remarkable advantage of LGCM within the structural equation modeling framework is that hypotheses are tested with latent (i.e., unobserved) variables estimated independently of the effects of measurement error in the observed variables (Bollen & Noble, 2011; Curran et al., 2010). Observed data were thus standardized (i.e., mean-centered and scaled by the standard deviation of each exogenous variable; in the case of latent variable indicators, the mean and standard deviation of the last time point was used for the standardization) prior to model fit. The latent intercept corresponds to the last follow-up. Accordingly, the latent slope weights used were −2, −1, and 0, for baseline, first, and second follow-up, respectively. The pattern of data missingness was examined for indications of data missing at random and further tested with Little’s (R. J. A. Little, 1988) test as implemented in the *mcar_test* function of the *naniar* package in R. Multivariate normality was verified through Henze-Zirkler’s multivariate normality test as implemented in the *MVN* R package, which also includes Anderson-Darling’s univariate normality test. The model was thus estimated with full information maximum likelihood (FIML), a method that allows all available data to be used for parameter estimation under the assumptions of data missing at random and multivariate normality (Enders & Bandalos, 2001). When assumptions are met, FIML estimation is the recommended practice for good external validity in longitudinal analysis with attrition (e.g., Little et al., 2014). The LGCM was estimated using the ‘lavaan’ package (v. 0.6-9; Rosseel, 2012) (https://lavaan.ugent.be/) in R. Model parameter estimates are reported as standardized.

### Longitudinal measurement invariance

Longitudinal measurement invariance was confirmed by no change in CFI and RMSEA when imposing model constraints to be equal across time points, and by non-significant results in the Chi-squared difference test between three nested measurement models that gradually increased the constraints imposed on them (see Supplementary Table S4). Multivariate (HZ = 0.84, *p*-value = 0.547) and univariate (Anderson-Darling statistic range = 0.22 – 0.69, *p*-values > 0.062) normality was confirmed for baseline age, baseline SCD, CON-FC for each time point, and CON-SC averaged across time points.

### Conditional CON-FC LGCM

A conditional LGCM tested whether SCD (as indexed by the inverse score of the MFQ) at baseline predicts CON-FC (averaged across all CON-FC ROIs) at the last follow-up (latent intercept) and/or the rate of change towards the last follow-up of CON-FC (latent slope) (second upper square from left to right and corresponding arrows in Fig. 1). CON-FC was obtained by averaging over all ROIs to have a single measure for the entire network, given that no differential effects of SCD were expected for specific ROI pairs. The latent intercept of CON-FC was centered at the last time point (i.e., as denoted by the −2.00, −1.00, 0.00 factor loadings, from left to right, next to the dotted line arrows in Fig. 3) – similar to the approach used in Daugherty and Raz (2016). The interaction between CON-SC and SCD was included to test the moderation of CON-SC on the effect of SCD on CON-FC as a predictor of the latent intercept of CON-FC (leftmost upper square in Fig. 1). As CON-SC was obtained by averaging across all time points, its direct effect was also included in the model, i.e., by pointing towards the latent slope of CON-FC (rightmost upper square in Fig. 1). For completeness, an alternative model was tested in which the moderation and direct effects of CON-SC were swapped with respect to the latent intercept and slope of CON-FC (i.e., with the moderation pointing to the slope and direct effect pointing to the intercept). Effects were considered significant at a Bonferroni-corrected α-level = 0.025 (i.e., to account for the two models tested). Age at baseline was used as a (time-invariant) control covariate, and all observed measures were centered at 0 with a standard deviation of 1.

**Fig 1.**
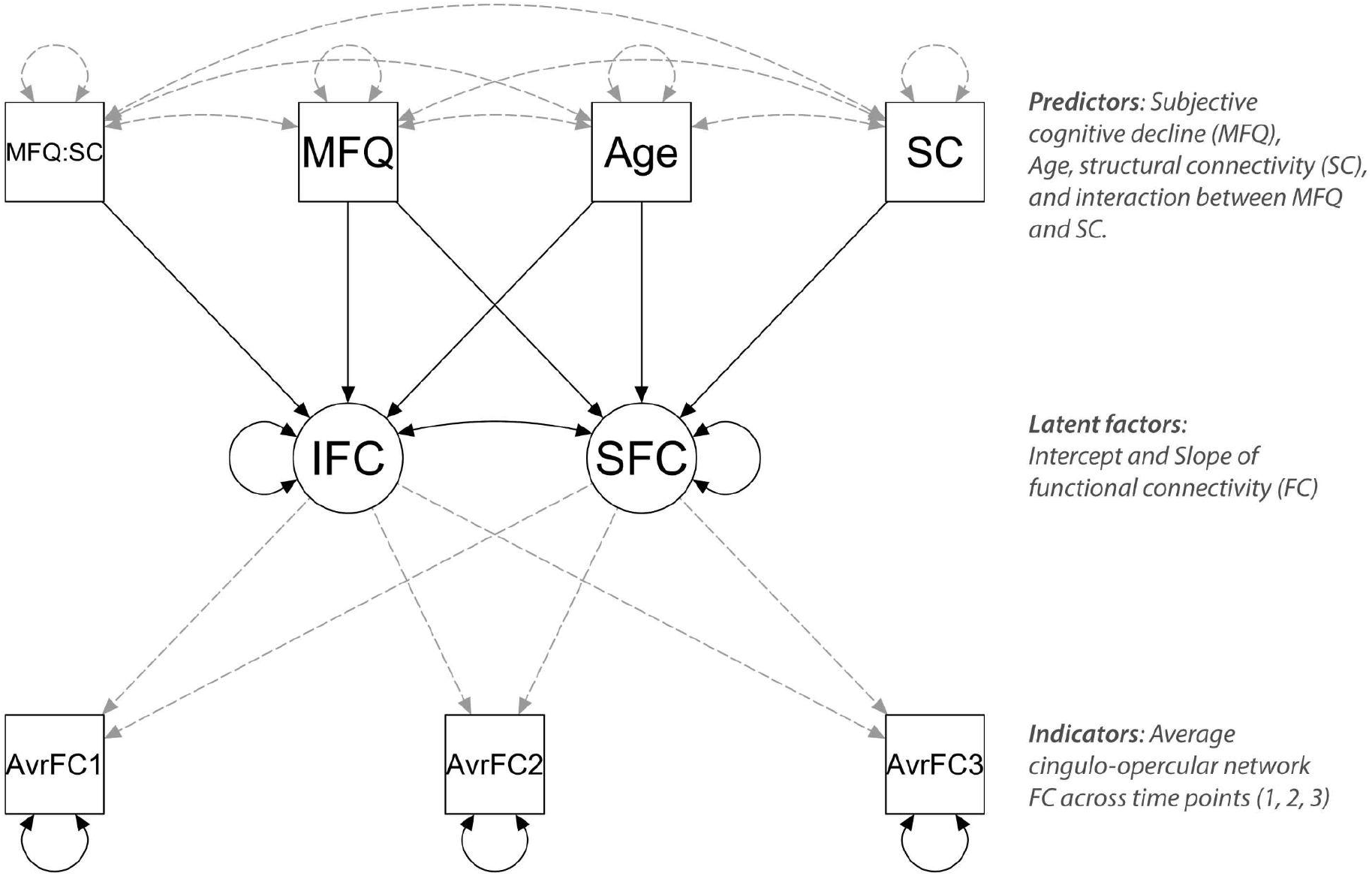
Conditional latent growth curve model path diagram. Model to test the effect of subjective cognitive decline, measured with the Memory Functioning Questionnaire (MFQ), on the latent intercept (‘IFC’) and slope (‘SFC’) of the functional connectivity averaged across regions of interest of the cingulo-opercular network (‘AvrFC’) for each measurement occasion. The model also includes the effect of structural connectivity (‘SC’) of the forceps minor and the interaction between MFQ and SC (‘MFQ:SC’). The directional effects of ‘SC’ and ‘MFQ:SC’ were also tested on the latent intercept and slope, respectively. Exogenous or predictor variables are on the top. Black arrows indicate that the effects were estimated, whereas gray, dotted arrows indicate that the effects were fixed. Squares represent manifest variables, while circles latent variables. Straight arrows indicate the directionality of a direct effect. Curved, double-headed arrows represent correlations or residual variances.

### Model evaluation

As there is ‘error’ (or unexplained variance) inherently associated with the measurement required to estimate CON-FC on each occasion, we ensured good consistency of measurement across study occasions to support the validity of the latent variables. Accordingly, invariance over time was assessed in the measurement model (latent variables and their three indicators; middle and lower part of Fig. 1) and ensured in the structural model (Fig. 1) by estimating variances within a latent construct and constraining them to be equal across time (van de Schoot et al., 2012).

Model fit was determined by examining the Chi-square (χ^2^) statistic (a non-significant value for acceptance), the comparative fit index (CFI ≥ .95 for acceptance), the root mean square error of approximation (RMSEA < .08 for acceptance), and the standardized root mean square residual (SRMR ≤ .08 for acceptance) (Schreiber et al., 2006).

### Data availability

The structured data and scripts used to generate these results are openly available and can be downloaded from https://osf.io/zaef2/.

## Results

### Behavioral and neuropsychological measures

All behavioral, and neuropsychological measures across the three time points are listed in Table 2. Linear mixed model analyses revealed that depressive symptoms increased and conscientiousness scores decreased across time points and with higher baseline SCD (see Supplementary Tables S2 and S3). There was also a time point × SCD interaction for both depressive symptoms and conscientiousness. For both depressive symptoms and conscientiousness, this interaction indicated a reduced association with SCD at the second follow-up. For depressive symptoms, the association became less positive with time, whereas for conscientiousness, the association became less negative with time. No other behavioral or neuropsychological measure varied across time points or with the degree of SCD. Age and SCD at baseline were not correlated (*r*_68_ = 0.14, 95% CI [-0.11, 0.36], *p* = 0.269).

### CON structural and functional connectivity

The ROIs from which the respective connectivity values were extracted for the LGCM are shown in Fig. 2A (CON-SC) and Fig. 2C (CON-FC). The line plots in Fig. 2B (CON-SC) and Fig. 2D (CON-FC) further illustrate the mean and variability in those values in the present sample across time points. CON-SC (averaged across time points) was negatively correlated with age at baseline (*r*_67_ = −0.35, 95% CI [-0.55, −0.12], *p* = 0.003). CON-SC tended to correlate negatively with baseline SCD (*r*_66_ = −0.22, 95% CI [-0.44, 0.02], *p* = 0.072). CON-SC did not differ across levels of education [*F*(4, 61) = 1.79, *p* = 0.143] or between males and females [*F*(1, 65) = 2.66, *p* = 0.108].

**Fig 2.**
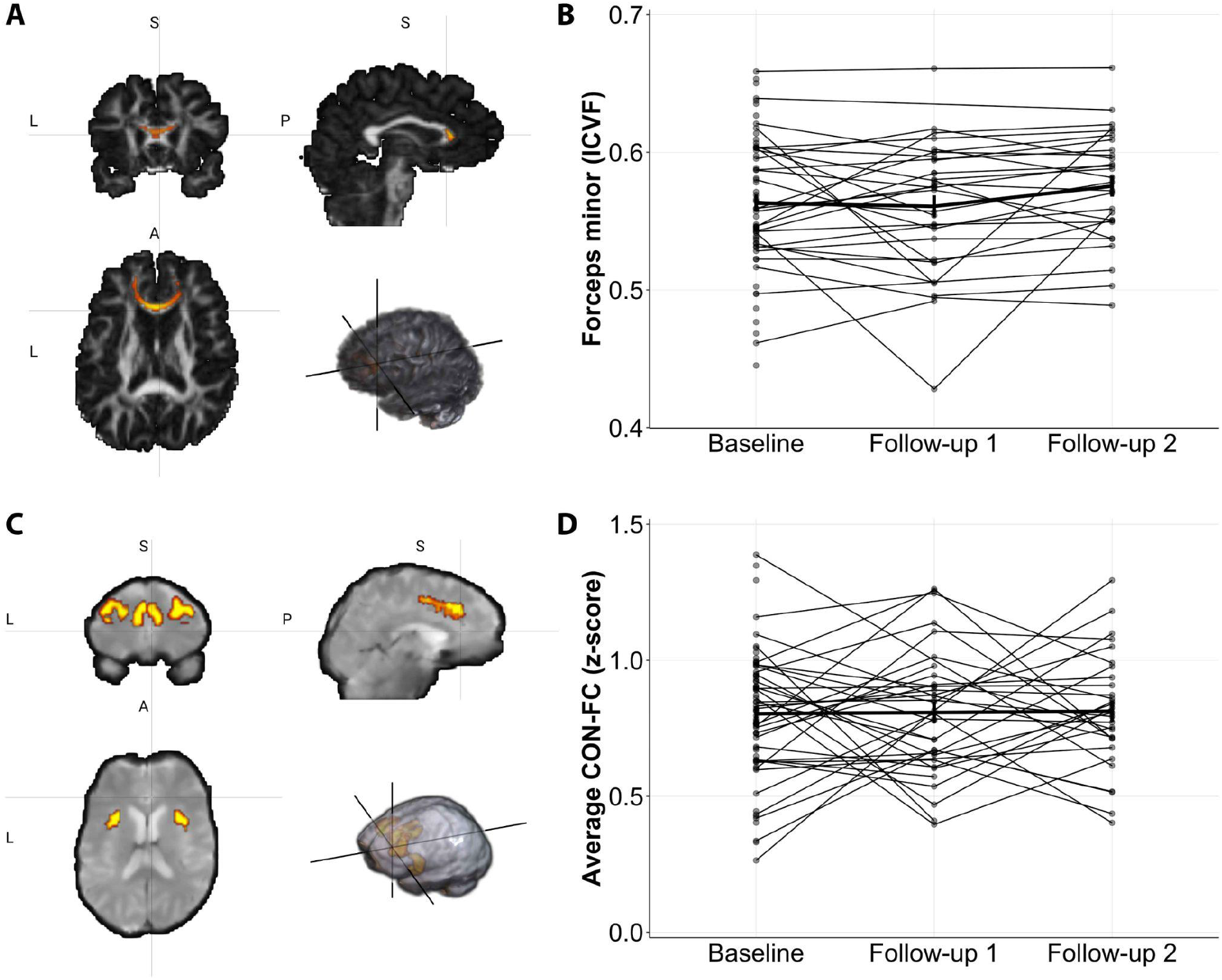
Individual and mean values of structural (SC) and functional (FC) connectivity of the cingulo-opercular network (CON). (A) Reconstructed forceps minor map in an exemplary participant. Neurite density (i.e., intracellular volume fraction, ICVF) values were obtained from this region of interest (ROI) for each participant and each time point as our measure of SC. (B) Individual and mean (thicker line) ICVF values of the forceps minor for each timepoint. The *y*-axis scale was chosen to facilitate the visualization of individual lines. (C) ROIs that composed the CON. To measure FC, correlations were computed between these ROIs (see Table 3) and an average was obtained for each participant and each time point. (D) Individual and mean (thicker line) FC values of the CON for each time point. The differing voxel intensity values in the ROIs were used to compute weighted averages of neurite density and time series in A and C, respectively. Error bars represent the standard errors of the respective means.

### CON-SC modifies the effect of SCD on CON-FC

The conditional LGCM of CON-FC had adequate fit (χ^2^ (9, *n* = 69) = 8.52, *p* = 0.483, CFI = 1.0, RMSEA = 0.0, and SRMR = 0.097). On average, SCD (as indexed by a higher MFQ) did not predict CON-FC after three years (latent intercept: β = −0.36, b = −0.19, Standard Error, SE = 0.15, 95% CI [-0.49, 0.12], *p* = 0.211) or its rate of change (latent slope: β = −0.53, b = −0.17, SE = 0.10, 95% CI [-0.38, 0.03], *p* = 0.093), contrary to what was hypothesized. However, CON-SC (averaged across all time points) moderated the relation between baseline SCD and CON-FC after three years (β = 0.49, b = 0.21, SE = 0.09, 95% CI [0.03, 0.39], *p* = 0.021; Fig. 3A), indicating that high baseline SCD predicts lower CON-FC after 3 years *given* a relatively low level of CON-SC throughout the 3-year time period (Fig. 3B). Neither the effect of CON-SC on the latent slope of CON-FC (95% CI [-0.19, 0.08], *p* = 0.399) nor the effect of age on the latent intercept (95% CI [-0.33, 0.23], *p* = 0.711) or on the slope of CON-FC (95% CI [-0.17, 0.22], *p* = 0.812) were significant.

**Fig 3.**
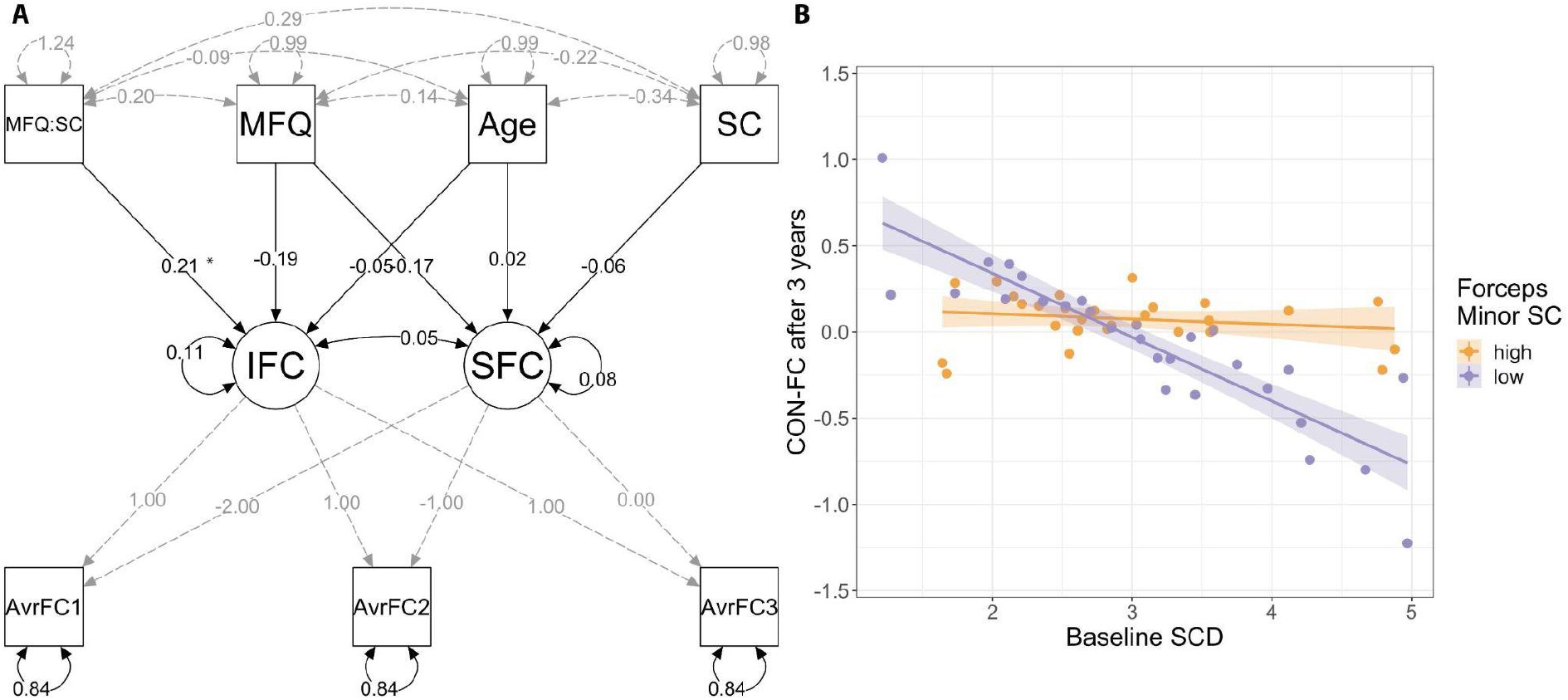
Conditional LGCM of CON-FC. (A) The effects of SCD (indexed by the inverted scores in the MFQ) on the latent intercept (‘IFC’) and slope (‘SFC’) of CON-FC were tested as well as the moderation of CON-SC on the relationship between MFQ and IFC (‘MFQ:SC’; arrow from the left-most top square). The moderation effect is illustrated in (B). (B) CON-SC, represented by the forceps minor neurite density, moderated the relationship between baseline SCD and CON-FC after 3 years. Note that the division into high and low CON-SC is for illustrative purposes only; the analysis was performed using CON-SC as a continuous variable. Unstandardized estimates are shown. * *p* < 0.025.

For completeness, an alternative model was tested in which the moderation and direct effects of CON-SC were swapped with respect to the latent intercept and slope of CON-FC – that is, CON-SC may plausibly moderate the rate of change (i.e., slope) of CON-FC. The model had adequate fit (χ^2^ (9, *n* = 69) = 6.61, *p* = 0.678, CFI = 1.0, RMSEA = 0.0, and SRMR = 0.079), and there was no evidence to suggest moderation; therefore, the alternative hypothesis of CON-SC moderation of the relationship between baseline SCD and the rate of change in CON-FC was rejected (95% CI [-0.20, 0.02], *p* = 0.111) (see Supplementary Fig. S3). A marginal, nominally significant effect was observed for CON-SC on the latent intercept of CON-FC that did not survive family-wise error correction (β = 0.76, b = 0.24, SE = 0.12, 95% CI [0.01, 0.47], *p* = 0.036, *ns* at the Bonferroni-corrected α-level). As with the previous model, neither SCD (intercept: 95% CI [-0.45, 0.13], *p* = 0.278; slope: 95% CI [-0.36, 0.03], *p* = 0.092) nor age (intercept: 95% CI [-0.25, 0.29], *p* = 0.882; slope: 95% CI [-0.14, 0.23], *p* = 0.662) predicted the latent intercept or slope of CON-FC.

## Discussion

In the present study, we tested whether baseline SCD predicts a decrease in CON-FC over three years and whether the decrease in CON-FC is more pronounced in older adults with relatively lower neurite density in the forceps minor (denoted as ‘CON-SC’, for simplicity). We used LGCM to study individual differences in CON functional and structural connectivity in a sample of healthy older adults. We found that greater SCD predicts lower CON-FC after 3 years in individuals with relatively low neurite density in the forceps minor. However, greater SCD did not directly predict lower CON-FC after three years or a greater rate of change in CON-FC over this period. Together, these results both indicate that CON-FC alterations in SCD are dependent upon the forceps minor neurite density and support the complex effect of SCD on FC.

Our main finding is that the neurite density in the forceps minor moderated the effect of SCD on CON-FC. SCD is a behavioral phenotype with multiple causes and trajectories in relation to cognitive function (Jessen et al., 2020; Rabin et al., 2017). Our finding thus indicates that one trajectory of SCD leading to functional brain changes such as decreased CON-FC depends on the neurite density of, at least, one major frontal white-matter tract, the forceps minor. This is not to say that other major (or minor) white-matter tracts are of lesser importance or no relevance for the CON-FC, given that an exhaustive exploration of CON-SC was beyond the scope of the present study (see *Supplementary Material).* Nevertheless, our finding does indicate that neurite density in the forceps minor might be a potential candidate biomarker for functional brain changes in individuals with SCD. The functional relevance of the frontal white matter, including the genu of the corpus callosum, has been implied by the cross-sectional association between fractional anisotropy in the frontal white matter and processing speed in healthy older adults (Hong et al., 2015). Future longitudinal studies could help elucidate whether decreased visual processing speed, one of the cognitive functions supported by the CON-FC (Ruiz-Rizzo et al.,2019), depends on the neurite density in, specifically, the forceps minor in SCD.

Our analyses revealed that the neurite density of the forceps minor was negatively correlated with age, in line with previous reports for fractional anisotropy and mean diffusivity (de Groot et al., 2015), voxelwise neurite density (Merluzzi et al.,2016), and overall diffusion metrics from atlas-based tracts of interest (Raghavan et al., 2021). Neither education nor sex was associated with the neurite density of the forceps minor in the present sample, also in line with previous reports (e.g., Raghavan et al., 2021). The association observed with age helps us better understand the moderating role of CON-SC in the effect of SCD on CON-FC, found in the present study. Specifically, the effect of SCD on CON-FC appears to be stronger for older adults who are more sensitive to age effects on CON-SC. Aging is associated with a high load of white matter hyperintensities, which tend to accumulate, mostly, in the frontal lobes (Raz et al., 2007). White matter hyperintensities are indicative of small vessel disease (Wardlaw et al., 2015) and have been shown to be associated with lower neurite density of, mainly, corpus callosum (i.e., including the forceps minor) and association (e.g., superior longitudinal fasciculus) fibers (Raghavan et al., 2021). Accordingly, it appears plausible that vascular risk factors or particular age-related vascular damage in frontal regions might contribute to a neurite density reduction in the forceps minor. Although education level did not particularly relate to CON-SC in the present sample, it is possible that other indicators of brain reserve or maintenance (e.g., cognitive, social, or physical activity) might influence vascular risk factors or overall neurite density reduction in the white matter. In line with previous studies (e.g.,Wang et al., 2012; Wen et al., 2019), we observed a negative but non-significant correlation between SCD and CON-SC. Larger sample sizes or a longer follow-up period in future studies might help determine whether there indeed is a direct association between SCD and CON-SC. Regardless, our results imply that CON-SC might allow identifying individuals with SCD in whom prevention strategies (e.g., aimed at reducing vascular risk factors, such as through dietary or physical activity interventions) may counteract functional brain decreases.

The LGCM approach used in the present study allowed us to separate the effect of the interaction between CON-SC and SCD on CON-FC at a particular time throughout the 3-year measurement period. Contrary to what was observed for the latent intercept, CON-SC did not moderate the effect of SCD on the rate of change of CON-FC (latent slope). The observed tendency was an overall decrease with greater SCD, independently of the level of neurite density in the forceps minor. In previous studies, greater SCD has been found associated with a more pronounced cognitive decline in memory functions across follow-ups spanning substantially larger follow-up periods, i.e., ~11 years (Hohman et al., 2011). Similarly, rates of cognitive decline have been shown to differ between older adults with and without SCD only after 6 years from baseline (Koppara et al., 2015). Accordingly, and assuming a similar effect for FC, a measurement period > 3 years might help ascertain whether CON-SC does also modulate the rate of decrease of CON-FC or, more generally, whether SCD influences the rate of decrease of CON-FC. In this context, testing potential non-linear change trajectories is an interesting consideration for future studies. Overall, our results suggest that CON-SC is relevant for the prediction of the level of CON-FC after three years, in individuals with the subjective impression of a decline in their cognitive functions.

Our results should be interpreted taking some limitations into account. For example, our sample size is small for LGCM approaches, which work best with sample sizes ≥ 100(e.g., Curran et al., 2010). Nevertheless, our study can set the ground for future longitudinal studies that combine FC and SC measures in a theory-driven manner in the context of SCD. Additionally, both MRI-based functional and structural connectivity are indirect measures of connectivity, which, moreover, are not always closely related (i.e., there is no one-to-one correspondence between the two). Despite this limitation, our study showed that combining both can offer a more comprehensive understanding of the changes in brain organization that might occur with SCD. Depressive symptoms and conscientiousness scores increased with SCD and time in our sample, thereby raising the possibility that they can partly explain the associations found between SCD and brain connectivity. However, subclinical depressive symptoms (i.e., a score < 5 on the GDS; Bijl et al., 2006) and particular personality factors (such as high neuroticism or conscientiousness) often co-occur with SCD and might either result from SCD or share the same underlying cause as SCD (Jenkins et al., 2019; Jessen et al., 2020). Our sample was composed of mostly females across time points (more accentuated at baseline and at the first follow-up), which may limit the generalization of our findings and conclusions to male older adults. Finally, averaging CON-SC across all time points required making assumptions that could not be directly tested in the present study (e.g., measurement invariance, interchangeability of observations, and interpretation of the average) but that merit a deeper investigation in the future (e.g., with more complex approaches like parallel growth modeling, given sufficient sample size).

To conclude, we found that SCD predicts lower CON-FC over three years in individuals with relatively lower neurite density of the forceps minor, a major frontal white-matter tract. Advanced age can have an impact on the neurite density of the forceps minor. Together, these findings imply modifiable age-related factors that could help delay or mitigate both age and SCD-related effects on brain connectivity.

## Abbreviations

BOLD: blood oxygenation level-dependent signal
CON-FC: cingulo-opercular network functional connectivity
CON-SC: cingulo-opercular network structural connectivity
fMRI: functional magnetic resonance imaging
LGCM: Latent growth curve modeling
MFQ: memory functioning questionnaire
NODDI: Orientation Dispersion and Density Imaging
SCD: subjective cognitive decline

## Acknowledgments

This work was supported by the European Union’s Framework Programme for Research and Innovation Horizon 2020 (2014-2020) [Marie Skłodowska-Curie Grant Agreement No. 754388 (LMUResearchFellows)]; LMUexcellent [funded by the Federal Ministry of Education and Research (BMBF) and the Free State of Bavaria under the Excellence Strategy of the German Federal Government and the Länder]; and the Netherlands Organisation for Scientific Research [Veni grant: 016.136.072]

## CRediT author contribution

A.R.: Conceptualization, Data curation, Formal analysis, Methodology, Software, Visualization, Writing – original draft, Writing – review & editing; R.V.: Data curation, Formal analysis, Investigation, Resources, Software, Writing – review & editing; A.D.: Methodology, Writing – review & editing; K.F.: Supervision, Writing – review & editing; H.M.: Supervision, Writing – review & editing; J.D.: Conceptualization, Funding acquisition, Methodology, Supervision, Writing – review & editing.

## Footnotes

1 For each response *x* of the 33 factor items: (8 – *x*) / 33.

2 For each individual, a residualized functional image was obtained after the voxelwise regression of the demeaned and detrended whole-brain signal (a time course vector resulting from the average across all brain voxels) on the demeaned and detrended ICA-AROMA output.

