## Supplementary Material for "Subjective cognitive decline predicts lower cingulo-opercular network functional connectivity in individuals with lower neurite density in the forceps minor"

### Supplementary Methods

#### Cingulo-opercular network's functional connectivity

To confirm that the selected regions of interest (ROIs) overlap with previously reported cingulo-opercular network (CON) regions, we visually inspected the spatial overlap of these ROIs with those reported in Shirer et al. (2012) – which have guided previous functional connectivity (FC)-based tractography approaches (e.g., Figley et al., 2015). Further, to confirm that these ROIs did also resemble the CON, the voxelwise whole-brain FC was computed, for each participant and each time point, using FSL's *dual_regression* command and our ROIs as seeds. The individual FC maps were converted to MNI space with the *flirt* command using each dataset's 'example_func2standard.mat' transformation matrix. Next, these maps were used as input to compute group FC maps, separately for each ROI and time point, using FSL's *randomise*. Each of the group maps was compared to an independent CON map (Shirer et al., 2012) by determining their degree of spatial correlation (using the *fslcc* command).

Whole-brain voxelwise analyses confirmed that the CON-FC ROIs selected from the multimodal parcellation did form the CON across time points

(Supplementary Fig. S1). Visual inspection additionally confirmed that the selected ROIs did overlap with those derived from Shirer et al.'s network-based atlas (Supplementary Fig. S2).

### Supplementary Results

#### Other possibly relevant white-matter tracts: uncinate fasciculus, superior longitudinal fasciculus (parietal), and anterior thalamic radiation

*Post-hoc*, we explored whether other white-matter tracts connecting frontal regions with other brain regions could also moderate the association between baseline subjective cognitive decline (SCD) and CON-FC after three years. The models based on the neurite density (averaged across the three time-points) of the uncinate fasciculus and superior longitudinal fasciculus (each one averaged across hemispheres) could not be evaluated given that model indices indicated inadequate fit to the data [uncinate fasciculus: $\chi 2$ (9, n = 69) = 11.49, $p = 0.244$, CFI = 0.37, RMSEA = 0.06, and SRMR = 0.09; superior longitudinal fasciculus: $\chi 2$ (9, n = 69) = 14.21, $p = 0.115$, CFI = 0.46, RMSEA = 0.09, and SRMR = 0.10].

The model based on the anterior thalamic radiation, however, did show adequate model fit [$\chi 2$ (9, n = 69) = 8.67, $p = 0.468$, CFI = 1.00, RMSEA = 0.00, and SRMR = 0.09] and was, thus, assessed further. The structural model results were similar to the main results based on the forceps minor. More specifically, neurite density in the anterior thalamic radiation also moderated the association between baseline SCD and CON-FC after three years [latent intercept: $\beta = -0.46$, b = 0.17, Standard Error, SE = 0.08, 95% confidence interval, CI [0.01, 0.34], $p = 0.034$]. However, this result does not survive a multiple-comparison correction (e.g., at the level used for the forceps minor, i.e., 0.025).

**Supplementary Discussion**

We interpret the poor fit of the uncinate and superior longitudinal fasciculi as an indication that the moderation we observe for the forceps minor is not readily explained by overall white matter integrity – at least with regard to CON-FC, in particular. Moreover, the results based on the anterior thalamic radiation would also suggest a relative specificity for the forceps minor in the context of CON-FC and SCD. Nevertheless, given the hypothesis-driven analytical approach followed in our study, we cannot conclude that the forceps minor uniquely or exclusively supports CON-FC.

**Supplementary Figures**

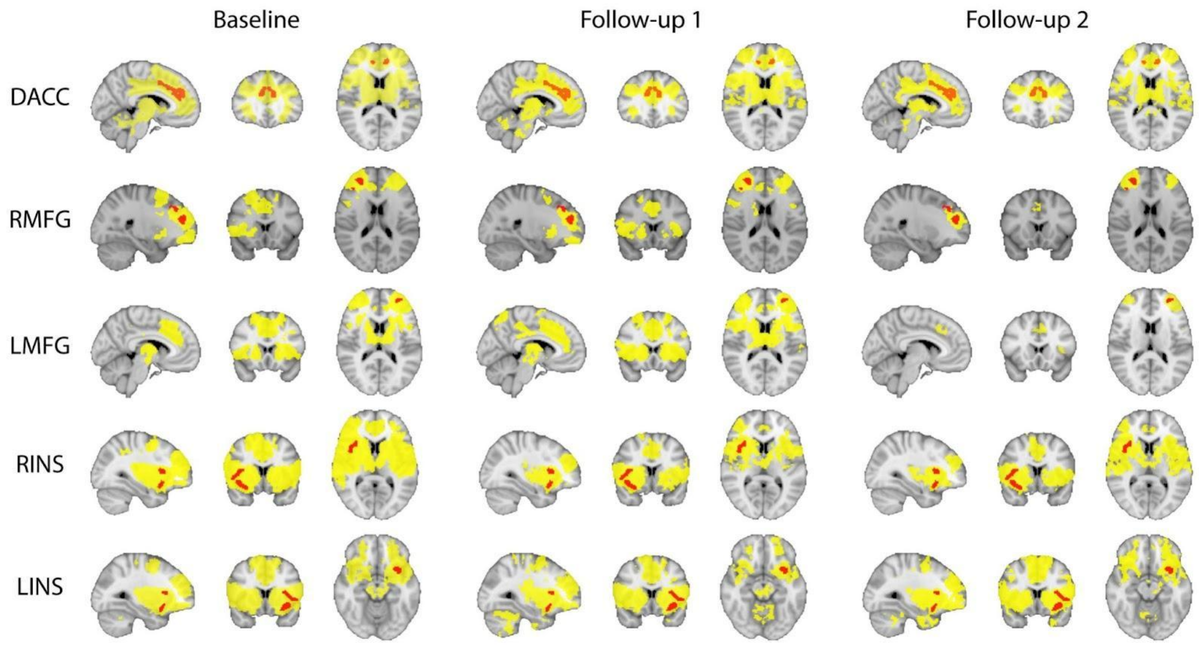


**Fig S1. Group functional connectivity (FC) maps of the cingulo-opercular network (CON) for seed region and time point.** Whole-brain, voxel-wise FC (in yellow) was verified for each region of interest (ROIs; overlays in red) and time point across individuals. Despite some variability across ROIs and time points, all maps include CON regions. DACC: dorsal anterior cingulate cortex; L/RINS: left/right insula. L/RMFG: left/right middle frontal gyrus. See main text for information on how these ROIs were selected and obtained. Maps are in radiological orientation (right is on the left).

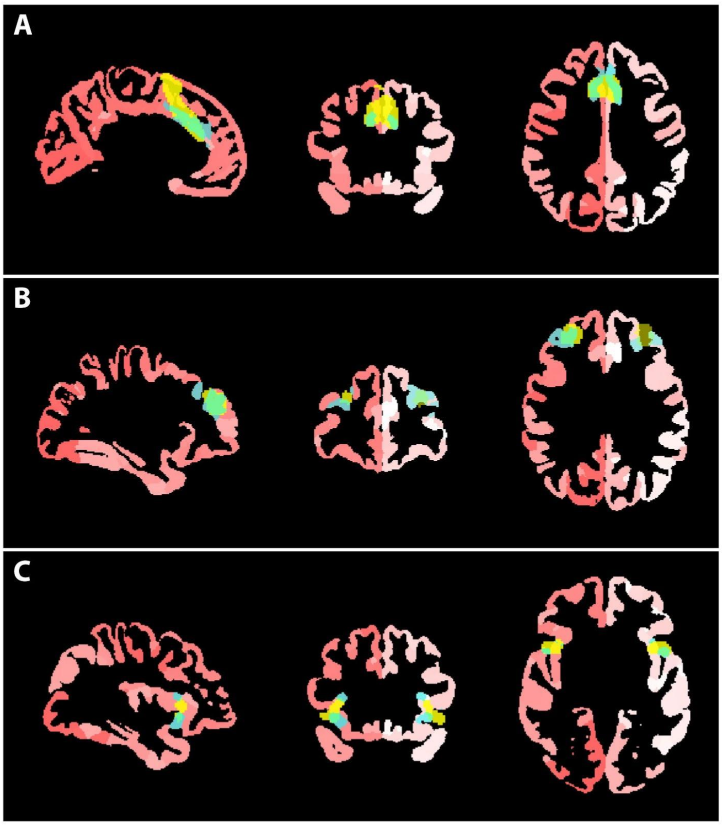


**Fig S2. Multimodal and functional regions of interest (ROIs) of the cingulo-opercular network (CON).** The selected multimodal ROIs (blue) (Glasser et al., 2016) overlap network-based functional ROIs (yellow) (Shirer et al., 2012). (A) anterior cingulate cortex, (B) left and right inferior/middle frontal gyrus, and (C) left and right anterior insula ROIs overlaid on the multimodal parcellation map of Glasser et al. (2016) in standard (Montreal Neurological Institute) space.

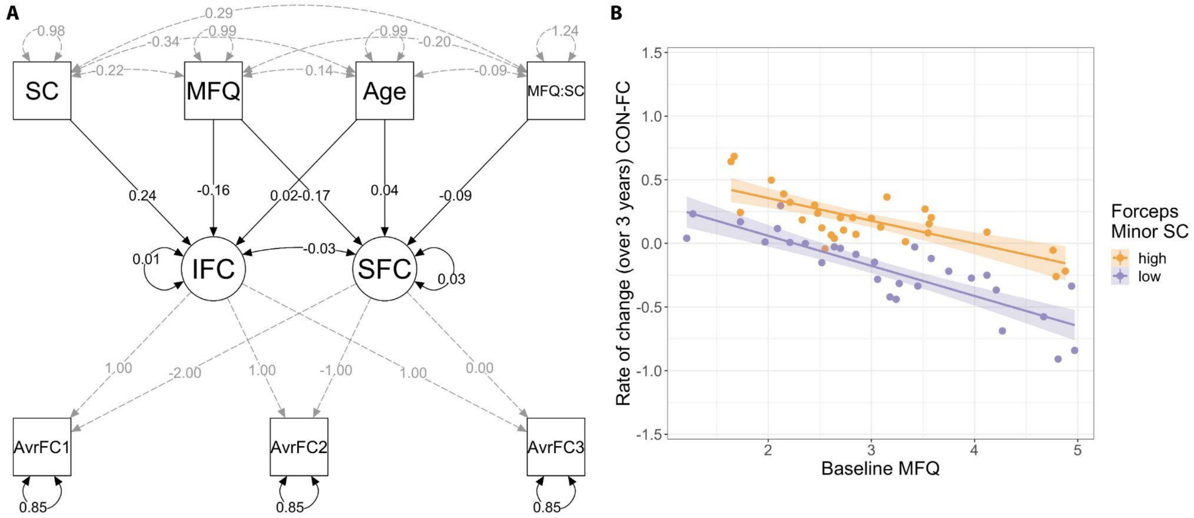


**Fig S3. Conditional LGCM of CON-FC with an alternative moderation of CON-SC.** (A) The effects of SCD ('MFQ') on the latent intercept ('IFC') and slope ('SFC') of CON-FC were tested as well as the moderation of CON-SC on the effect of MFQ on SFC. (B) CON-SC, represented by the forceps minor neurite density, did not moderate the effect of baseline SCD on the rate of change (SFC) of CON-FC.

### Supplementary Tables

**Supplementary Table S1.** Cross-correlation coefficients with the anterior salience network of Shirer et al. (2012)

| ROI | Baseline | Follow-up 1 | Follow-up 2 |
|---|---|---|---|
| ACC | 0.27 | 0.29 | 0.31 |
| LMFG | 0.35 | 0.29 | 0.30 |
| RMFG | 0.35 | 0.38 | 0.30 |
| LINS | 0.27 | 0.26 | 0.28 |
| RINS | 0.25 | 0.31 | 0.24 |

ACC: anterior cingulate cortex; L/RINS: left/right insula; L/RMFG: left/right middle frontal gyrus; ROI: region of interest. The whole-brain maps from which these cross-correlations were computed are shown in Fig S1.

**Supplementary Table S2.** Linear mixed model results for Geriatric Depression Scale (GDS) score

| **Fixed Effects** | ***b* (SE)** | **T-value** | **95% CI** | **Mean Squares, F-value** |
|---|---|---|---|---|
| Intercept | 0.91 (1.16) | 0.79 | [-1.35, 3.17] | |
| Time | 1.87 (0.84) | 2.23 | [0.23, 3.49] | 1.03, 0.34 |
| SCD (MFQ) | 0.97 (0.35) | 2.73 | [0.27, 1.67] | 11.60, 3.78 |
| Time × MFQ | -0.58 (0.27) | -2.12 | [-1.11, -0.05] | 13.82, 4.50 |
| **Random Effects** | **SD** | | | |
| Participant | 3.41 | | | |
| Residual | 1.75 | | | |

Formula: GDS score = Intercept + $b_1$(Time) + $b_2$(SCD) + $b_3$(Time * SCD) + Participant + Residual
Measurement occasion (Time) was coded starting with 0 at baseline. *Abbreviations*: CI: confidence interval; MFQ: memory functioning questionnaire; SCD: Subjective cognitive decline; SD: standard deviation; SE: standard error.

**Supplementary Table S3.** Linear mixed model results for Conscientiousness personality score

| **Fixed Effects** | ***b*** **(SE)** | **T-value** | **95% CI** | **Mean Squares, F-value** |
|---|---|---|---|---|
| Intercept | 43.87 (1.57) | 27.80 | [40.72, 47.00] | |
| Time | -3.53 (1.20) | -2.93 | [-5.86, -1.16] | 13.14, 2.03 |
| SCD (MFQ) | -2.18 (0.49) | -4.44 | [-3.17, -1.20] | 86.22, 13.30 |
| Time × MFQ | 1.02 (0.39) | 2.58 | [0.24, 1.78] | 43.33, 6.68 |
| **Random Effects** | **SD** | | | |
| Participant | 4.10 | | | |
| Residual | 2.55 | | | |

Formula: Conscientiousness score = Intercept + $b_1$(Time) + $b_2$(SCD) + $b_3$(Time * SCD) + Participant + Residual

**Table S4.** Longitudinal measurement invariance

| **Model** | **CFI** | $\chi^2$ **(*df*)** | **AIC** | **BIC** | **ΔCFI** | $\Delta\chi^2$ **(Δ*df*)** | ***p*-value** |
|---|---|---|---|---|---|---|---|
| Configural | 1.0 | 0.431 (1) | 399.75 | 417.62 | - | - | - |
| Scalar | 1.0 | 0.522 (3) | 395.84 | 409.25 | 0.0 | 0.091 (2) | 0.955 |
| Residual | 1.0 | 0.592 (5) | 391.91 | 400.85 | 0.0 | 0.070 (2) | 0.966 |

Measurement models: Configural: unconstrained measurement model with free estimation of item intercepts; Scalar: as Configural, but with item intercepts constrained to be equal between the three time points; Residual: as Scalar, but with item residuals constrained to be equal between the three time points. AIC: Akaike information criterion; BIC: Bayesian information criterion; df: degrees of freedom.
